# A Generalized Similarity Metric for Predicting Peptide Binding Affinity

**DOI:** 10.1101/654913

**Authors:** Jacob Rodriguez, Siddharth Rath, Jonathan Francis-Landau, Yekta Demirci, Burak Berk Üstündağ, Mehmet Sarikaya

**Affiliations:** GEMSEC, Genetically Engineered Materials Science and Engineering Center, University of Washington, Seattle, WA 98195, USA; Department of Materials Science and Engineering, University of Washington, Seattle, WA 98195, USA; Molecular Engineering and Science Institute, University of Washington, Seattle, WA 98195, USA; Department of Mathematics, University of Washington, Seattle, WA 98195, USA; Department of Electrical and Electronics Engineering, Middle East Technical University; Department of Computer and Informatics Engineering, Istanbul Technical University; Department of Chemical Engineering, University of Washington, Seattle, WA 98195, USA; Department of Oral Health Sciences, University of Washington, Seattle, WA 98195, USA

## Abstract

The ability to capture the relationship between similarity and functionality would enable the predictive design of peptide sequences for a wide range of implementations from developing new drugs to molecular scaffolds in tissue engineering and biomolecular building blocks in nanobiotechnology. Similarity matrices are widely used for detecting sequence homology but depend on the assumption that amino acid mutational frequencies reflected by each matrix are relevant to the system in which they are applied. Increasingly, neural networks and other statistical learning models solve problems related to functional prediction but avoid using known features to circumvent unconscious bias. We demonstrated an *iterative* alignment method that enhances predictive power of similarity matrices based on a similarity metric, the Total Similarity Score. A generalized method is provided for application to amino acid sequences from inorganic and organic systems by benchmarking it on the debut quartz-binder set and 3 peptide-protein sets from the Immune Epitope Database. Pearson and Spearman Rank Correlations show that by treating the gapless Total Similarity Score as a predictor of relative binding affinity, prediction of test data has a 0.5-0.7 Pearson and Spearman Rank correlation. with respect to size of the dataset. Since the benchmarks used herein are from a solid-binding peptide and a protein-peptide system, our proposed method could prove to be a highly effective general approach for establishing the predictive sequence-function relationships of among the peptides with different sequences and lengths in a wide range of biotechnology, nanomedicine and bioinformatics applications.

**Author Summary:** The significance of this work is to expand the applicability of a known metric for describing the function of tiny proteins also called peptides. The Total Similarity Score (TSS) can describe how ‘similar’ a peptide, or a group of peptides are to another group of sequences with a known or suspected function. A peptide/group of peptides will always have a high TSS if it contains the same or ‘similar’ amino acids in the same positions. This metric can therefore be used to select peptides for useful functions based purely on conserved amino acids in unknown positions. The greedy search algorithm used to learn how similar amino acids are to each other has been shown to be marginally effective in this larger dataset. Therefore, we argue that the TSS metric is a highly useful one for predicting peptide affinity but a different machine learning algorithm should be applied to make full use of it.

## Introduction

The rapid development of target-specific drugs relies on the development of high-throughput and accurate methods of modelling molecular structures. The biology, pharmacology and bioengineering communities are interested in building widely applicable methods founded in predictive design of molecules that have specificity for biological targets, analytes and biomarkers [1–4]. Small peptides (7 to 40 amino acids) have high potential as both therapeutics [5–7] and high-performance molecular building blocks [8–10] due their diversity of binding affinity both quantitatively and specifically across 2D- and nano-materials.

Towards more accurate and fast predictions of affinity or conformation that would enable high-throughput drug and targeting peptide design, among some of the best performing methodologies are stochastic models such as NetMHCpan-4.0 [11], DeepMHC [12] and MHCflurry [13]. These methods use little or no prior information about the peptides to ensure only random walk identifies relevant patterns. By avoiding physiochemical properties published in the literature, these models are subject to inconsistent predictions between test peptide sets even for the same protein target. Alignment-free neural networks models have shown substantial success in predicting the binding affinity of the Immune Epitope Database (IEDB, www.iedb.org) datasets [12,14]. To avoid overfitting, they require hundreds of thousands of sequences and are not optimized for gaps in the binding domains [15,16].

The current state of the art in modelling tools, e.g., molecular dynamics (MD), molecular mechanics (MM), and Monte Carlo (MC) based methods, predict overall conformation from which binding energies may be calculated [9]. These approaches utilize knowledge-based force fields [17,18] and energy minimization techniques to sample the most probable structures [19]. Though solving conformational structures will likely enable the most accurate predictions of peptide function, to date structural information is avoided in models requiring large amounts of data. This is mostly due to the large computational cost associated with calculating molecular structures of these large molecules, which is a barrier to the development of both highly complex neural networks and current MD/MC-based methods. The deeper networks rely less on learning in space constrained by verified physiochemical trends and more on the number of parameters and computational power. Less complex and more interpretable models integrate known patterns while leaving space for optimization methods to learn unknown patterns in the sequences.

Current alignment-based methods for high-throughput prediction functionality of amino acid sequence information can be separated into two groups; pairwise [20–22] and multiple sequence [16,23,24]. In general, pairwise alignment is ideal for shorter sequences due to its higher computational cost per amino acid and is widely accepted to be the optimal alignment [25]. Multiple sequence alignment is considered more appropriate for longer sequences with suspected consensus domains. In both methods Point Accepted Mutation (PAM) and Blocks Substitution Matrix (BLOSUM) matrices are still the most widely used, and there are permutations of these matrices to serve more specific tasks [17,26,27]. Overall, the limitations of PAM and BLOSUM provided inspiration and guidance for generating matrices with increased accuracy based on larger and more complete datasets [11,28–30]. Matrices such the PMBEC [27], have been generated based on the two models that produce a minor increase in performance but ultimately are vulnerable to the same factors as their predecessors [11]. In 2008, for example, a miscalculation was discovered in the clustering protocol of the BLOSUM matrix [31]. Despite extensive characterization of the mistake, BLOSUM is still the standard for one of the largest alignment-capable databases available to date, BLAST [11].

In contrast with PAM, BLOSUM, PMBEC [27] and the SAUSAGE Force Field Matrix [17], the novel OCSimM and 8 property group-derived matrices (A-RMat) were calculated from 527 physiochemical properties of amino acids [32,33]. AAindex is a vast resource of high-quality amino acid properties collected from literature dating from the mid-sixties to today [32,33]. Typically, either variable reduction methods (Principal Component Analysis [6,34] or Factor Analysis [35]) or heuristic selection is performed to shrink the huge dataset of over 550 amino acid properties to obtain an interpretable solution. Variable reduction has significant advantages over a global analysis of heuristically grouped properties because human error cannot influence the potential relationships observed [35]. However, these methods still assume the relationship between high-specificity peptides and low-specificity peptides is described by physiochemical properties.

Previously, we have successfully used a matrix optimization method to a group of peptides that were categorized as strong, weak or medium binders based on their binding affinities to crystalline silica, quartz, using 40 sequences that were originally genetically selected using M13 phage display peptide library [25]. The novel metric called the Total Similarity Score (TSS_A-B_) describes the average Global Alignment score of all peptides from group-A to all of group-B [25]. The TSS score quantifies the similarity of a peptide to a functional peptide set (i.e. affinity for a solid material). By keeping random changes to a similarity matrix that increased the TSS_S-S_ (TSS of strong binders with strong binders), and decrease the TSS_S-W_ (TSS of strong binders with weak binders), a similarity matrix was obtained that could predict the semi-quantitative affinity of quartz-binding peptides with 70-80% success. Despite its high predictive power, TSS has never been applied to MHC data. Using the MHC data, here we demonstrate its implementation that strongly suggests that TSS could be a predictive method for establishing sequence-function relationships in a variety of large sequence-based data sets.

The reliable prediction of peptide binding affinity has already led to ground-breaking advances in oral health science and will continue to do so in areas requiring a well-described soft interface between peptides and solid-state inorganic materials [5,10,36]. Though affinity prediction is not the most descriptive or important characterization of peptides, understanding the relationship among solid-binding peptides has led to many technologies such as sensors with high sensitivity, [5] assemblers in nanotechnology, and tiny enzymes in biomineralization [37].

## Materials & Methods

Iterative Alignment (IA) creates a scoring matrix that provides scores correlating with the positional composition of a peptide when compared to a weak and a strong binding set. When a sequence of interest has high similarity to these strong binders and low similarity to the weak binders, the sequence was given a higher TSS_Seq-S_ (TSS of one sequence to strong binders) and lower TSS_Seq-W_ (TSS of one sequence to weak binders). Training the similarity matrix was done by increasing the differences in TSS to strong binders for two binding affinity classes, strong and weak.

First, peptides were sorted by their affinity values shown in Fig 1A. The generated trend was characterized by the positive correlation TSS_Seq-S_ with binding affinity visualized in the lower bar chart of Fig 1B. Once training was finished, we calculated the TSS to strong binders a final time for all peptides in the set. The results from the trained randomly initialized matrix (RandM) are shown in the scatterplot in Fig 1C. Next, several methods are used to measure correlation of TSS_Seq-S_ with the experimental affinity including Pearson and Spearman Rank correlation, Root Mean Square Error and a binary classification scheme (binder/nonbinder prediction). A sample of these results are demonstrated in Fig 1D.

**Fig 1.**
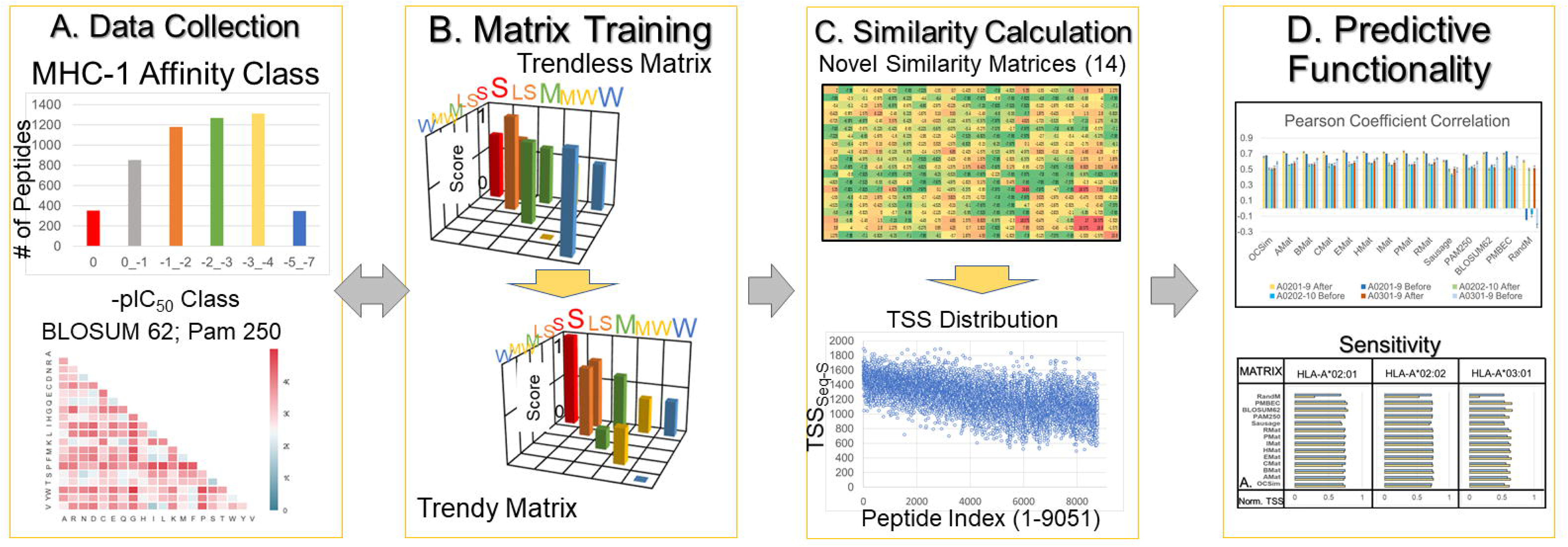
Schematic of Iterative Alignment Procedure. The iterative alignment procedure is executed in four separate steps as show in the flow chart, that include: (A) Classification of MHC-I binding peptides from the IEDB and the resultant matrices from AAindex; (B) Training the randomly initialized matrix (RandM) which, before training, was uncapable to demonstrate the trend of decreasing cross-similarity, but after training it becomes prominent indicating the successful integration of the information; (C) Demonstration the total similarity score of the full allele set with respect to the strongest determined binders of *HLA-A*02:01* (TSS _*HLA-A*02:01*_-S) for trained RandM. Calculations were performed for all matrices before and after training; (D) Showcase correlation and accuracy measurements (see details in the text and figures below).

### Data Collection

Peptide sequences with affinity for HLA alleles were obtained from the Immune Epitope Database (www.iedb.org), a common source of training and benchmark data for predictive models of peptide function [14]. Quartz binders and the Quartz I matrix were provided by GEMSEC at the MSE Department of the University of Washington [25]. The Amino Acid Index (AAindex) is a large database of amino acid properties that were used to calculate the cluster matrices (A-RMat) [32,33]. Within the site, similarity matrices calculated by various studies are also provided, and it was from here that the SAUSAGE force-field matrix was also chosen [17]. The PMBEC scoring-matrix [27] was included as it was derived directly from binding affinity data from MHC-I. In general, the matrices chosen are a diverse subsection of the types of information used to describe differences between amino acids and therefore were an appropriate selection for yielding conclusions about how the seed matrix would affect the overall result.

### Novel Matrix Calculation

To explore the possibility that certain properties, e.g., hydrophobicity, electrical properties, amino acid composition etc., may make better seed matrices, 9 similarity matrices were calculated based on clusters optimized by Saha *et. al* [38]. After grouping properties by alpha-helix or beta-sheet propensities, composition, electrical, hydrophobic, and intrinsic characteristics, residue propensity, and physicochemical properties, we performed Principal Component Analysis (PCA) on each group and all groups combined. Using a Python library downloaded from scikit-learn.org [39], the principal components were calculated which were most representative of the internal variation of property subset. Because these principal components are orthogonal, Euclidean distance was the most appropriate for calculating the actual similarity matrix. By calculating the difference between the principal components of two amino acids, we were able to calculate nine (20 × 20) similarity matrices describing their quantitative physiochemical differences. These matrices will be referred to for the rest of the work as AMat (Alpha-helix propensity), BMat (Beta-sheet propensity), CMat (Composition), EMat (Electric), HMat (Hydrophobicity), IMat (Intrinsic propensity), PMat (Physiochemical), RMat (Residue propensity) and OCSim [Orthogonal Component Similarity matrix (all properties)].

### Code Implementation

The newest version of the algorithm was written in Python, using a gapless scoring method to calculate TSS scores. The gap calculation was excluded to rectify the issue created by the changing gap position in each sequence. Per peptide-peptide scoring operation (300 strong binders x 9000 peptides for *HLA-A*02:01*), per iteration (5000 due to randomly changing mutabilities) the gap is placed in one position. We suspected the gap made recognizing the consistent amino acids between iterations difficult. The debut implementation of the method [25] iteratively aligned less than 20 peptides per strong and weak binding group. The IEDB dataset being substantially larger (i.e., over 9000 peptides for the largest set) required the inclusion of more peptides per set in order to capture as many of the features pertaining to binding affinity as possible.

### Designation of Affinity Classes

The peptide sequences were first ordered by -pIC_50_, and then segregated into groups dependent on their affinity. For example, all peptides within the 3 chosen alleles (*HLA-A*02:01* [9-length], *HLA-A*02:02* [10-length] and *HLA-A*03:01* [9-length]) with a - pIC_50_ of 0 were named ‘strong’ (S) binders, creating 3 sets. The ‘weak’ (W) binders for the 9-length and 10-length sets were those with a -pIC_50_ of −5 to −7. From these, 80% of a strong or weak peptide list was randomly chosen as training sets to obtain cross-validation. To show the flexibility of the method, we chose several groups with differing distributions to demonstrate the improvements are still achieved when only partial data is available.

### Matrix Training

To begin, two lists of peptide sequences (at least 6 in each) must be obtained, one with higher ‘internal similarity’ and lower ‘internal similarity’. Critically, peptides with high binding affinity for the same material will also higher ‘internal similarity’ and those with low affinity will have low ‘internal similarity’ [25]. Internal similarity refers to the sum of Global Alignment (GA) scores of each peptide within a list to every other peptide within the same list. Global Alignment is commonly referred to as the Needleman-Wunsch algorithm or optimal alignment as it always obtains the optimum number and placement of gaps, resulting in the most similar domains being recognized and aligned when they are consensus [20]. It requires a similarity matrix to obtain scores between matches or mismatches of amino acids, and many of these have been calculated throughout the literature. For small peptides, it may not be the ideal alignment method considering their short length makes scoring the entire sequence important.

While guaranteeing the optimal alignment, Global Alignment is computationally very expensive and therefore impractical to apply to larger groups of sequences than those used in previous work [25] (10 - 20 sequences per strong and weak group). The updated method departs from the alignment methodology and scores peptides by their positional composition only, which is essentially the same score without the gap calculation. By greatly expanding the number of peptides used in the strong group, the significance of GA is reduced due to a wider range of domain types and locations being represented. In general, scoring with more peptides is just as beneficial as scoring a few with GA. Global Alignment expands the number of sequences a peptide will have consensus with; in a way making it appear as many peptides in the strong group. However, the domains being aligned and the values scoring the alignments are different from one iteration to another, resulting in a lack of consistent scoring between sequence domains. Therefore, we justify the departure from GA as both a necessity and a benefit to ensure the method runs within a practical time constraint.

The procedure for one iteration can be described in 6 steps (see Fig 2). After the affinity classes have been designated (Fig 2, Step 2), a seed similarity matrix is used to calculate TSS_S-S_ and TSS_S-W_ (same as internal but to separate group of peptides) similarity for each peptide (Fig 2, Step 3). External similarity is calculated by aligning the strong binders to each in the low internal similarity group. Within each list, the average is found and form the cost functions for IA, the Total Similarity Score Strong-Strong (TSS_S-S_) and Total Similarity Score Strong-Weak (TSS_S-W_), respectively. Mathematically, the expression for general TSS calculation is given by Equation (1) as

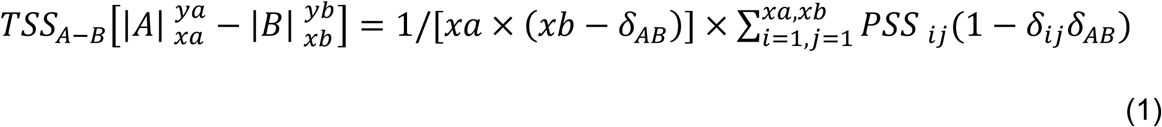

where, TSS_A-B_ is the Total Similarity Score (TSS) between peptide sets A and B, PSS_ij_ is the pairwise similarity score (PSS) between sequences i and j of sets A and B respectively, *xa* and *xb* are the total number of sequences in sets A and B, and *δ* is the Kronecker delta function (δ_ij_ = 1 if δ_ij_ = 0).

**Fig 2.**
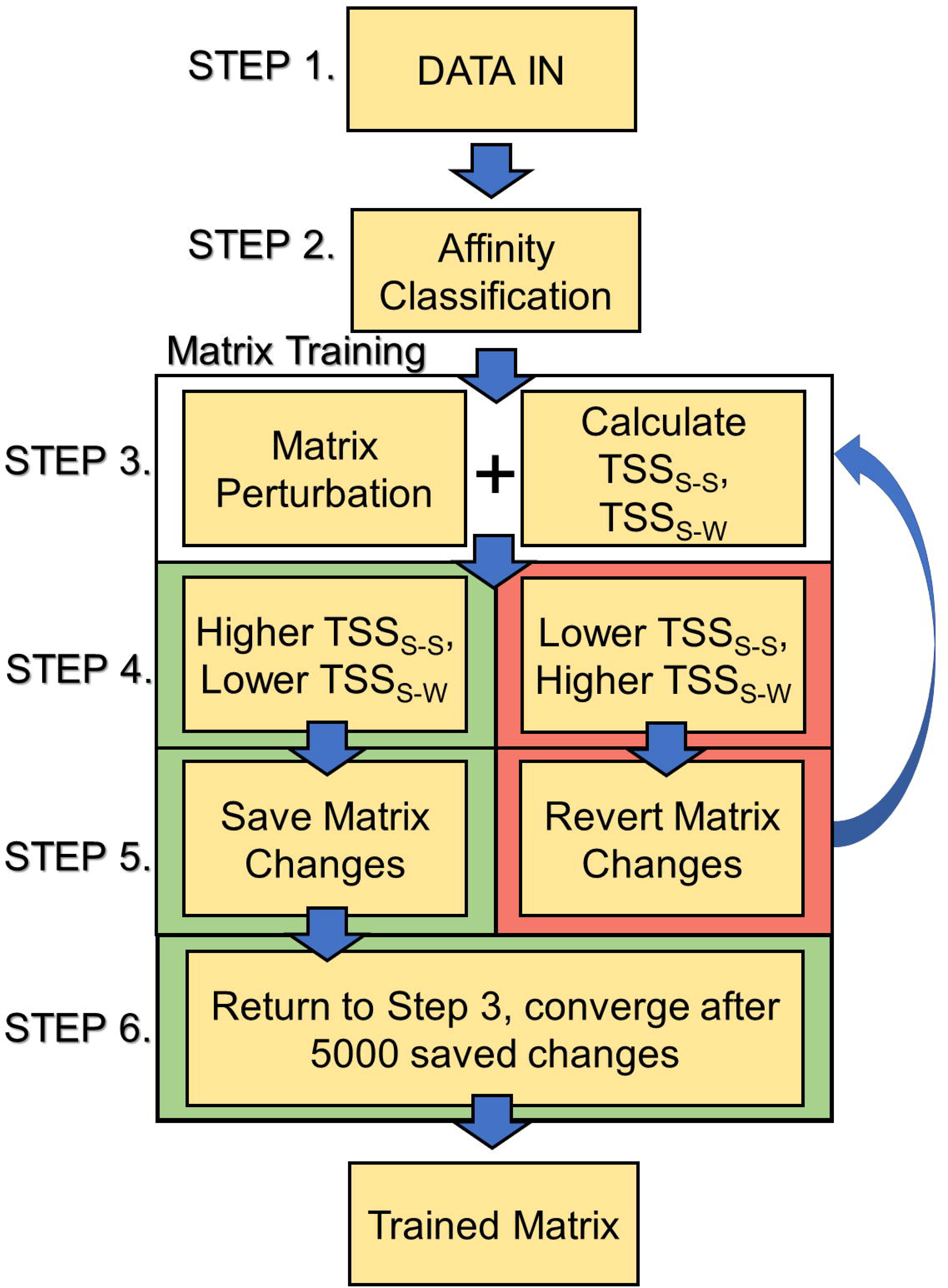
Schematics of matrix training procedure. Peptides were first downloaded and classified by their affinity. The similarity matrix is perturbed randomly and then TSS scores are calculated. Depending on the outcome, changes to the matrix were either saved or discarded. The matrix was considered ‘converged’ after 5000 beneficial changes total, or 5000 negative changes in a row, occur.

After the values of TSS_S-S_ and TSS_S-W_ have been calculated and saved for the first time, the similarity matrix is perturbed by making random changes (1-20) to the matrix values by either adding 1 or subtracting 1 (Fig 2, Step 3). Using the new matrix, TSS_S-S_ and TSS_S-W_ are calculated again and compared with the previous TSS (Fig 2, Step 4). A change to the matrix is considered beneficial if TSS_S-S,NEW_ is greater than TSS_S-S,OLD_ and TSS_S-W,NEW_ is less than TSS_S-W,OLD_. Beneficial changes are saved for the next round (Fig 2, Step 5). If the change is not beneficial, then the previous matrix (before mutation) is perturbed again and the process repeats (Fig 2, Step 5). The algorithm could continue indefinitely but we considered the matrix converged when over 5,000 iterations occurred without a beneficial change (Fig 2, Step 6).

### Benchmark with the Previous Work

To prove the updated methodology was up to par with the original implementation of the procedure, we obtained the Quartz I matrix and silica binding peptides used by Oren *et. al* [25] The same procedure was followed by mutating PAM250 and training on the same strong and weak groups. After training, IA converged on a matrix capable of predicting binding affinity with similar accuracy to the debut implementation [25]. Using a Pearson correlation of the external similarity to affinity of any silica binding peptide to the group of strong binders designated by [25], we calculated a 51% correlation with our matrix. Previous work obtained a 46% correlation with Quartz I, demonstrating the equivalent capabilities of the updated method. P-values for these correlations were less than 0.0005.

### Application to MHC Data

To test whether the modified methodology would perform on organic materials, we needed a set of peptides with affinity for a biological target. The IEDB provides high quality sequence data including binding affinities for multiple Major-Histocompatibility Complexes which provided a perfect opportunity to test performance [14]. By designating peptides with -pIC_50_ (negative logarithm of IC_50_) of 0 as strong-binders peptides and weak-binders having -pIC_50_ of −5 to −7 (Fig 2, Step 2) from three alleles (*HLA-A*02:01, HLA-A*03:01*, and *HLA-A*02:02*), we optimized 14 similarity matrices capable of ranking peptides by their binding affinities via their total similarity to strong binders. Matrices were optimized by iteratively perturbing a seed similarity matrix and keeping those changes which ultimately increased the self-similarity of the strong binders and cross-similarity of the strong with weak binders.

## Results/Discussions

### Cross-Similarity Analysis

Fig 3 shows the cross-similarity results of 5 subsets of peptides deemed Strong (S; -pIC_50_:0), Less Strong (LS; -pIC_50_:−1 to −2), Medium (M; - pIC_50_:−2 to −3), Medium Weak (MW; -pIC_50_:−3 to −4) and Weak (W; -pIC_50_:−5 to −7) based on their binding affinity to alleles MHC-I *HLA-A*02:01* for all matrices before and after training. Each set of bars per matrix was normalized by the largest value of both before and after results. In addition, these bars are the results of 5 average TSS subsets (80% randomly chosen from each affinity class). Previous work showed the TSS of a peptide with high similarity to the peptides that are strong binders of a solid-state material indicates that the peptide in question likely also has strong binding capability [25]. Therefore, the average TSS of peptides with an affinity for a protein should decrease with their experimental affinity. Fig 3 shows that before training (yellow bars) the trend is somewhat present but not very defined (Strong x Weak is comparable to Strong x Less-Weak) but after training (blue bars) the trend is very pronounced. For each matrix and across all three alleles (see S1 and S2 Figs) we observe average TSS when grouped by affinity class to strong binders correlated with experimental affinity. Most notably, the randomly initialized matrix RandM despite having no initial correlation was able to show the trend as definitively as the others after matrix training.

**Fig 3.**
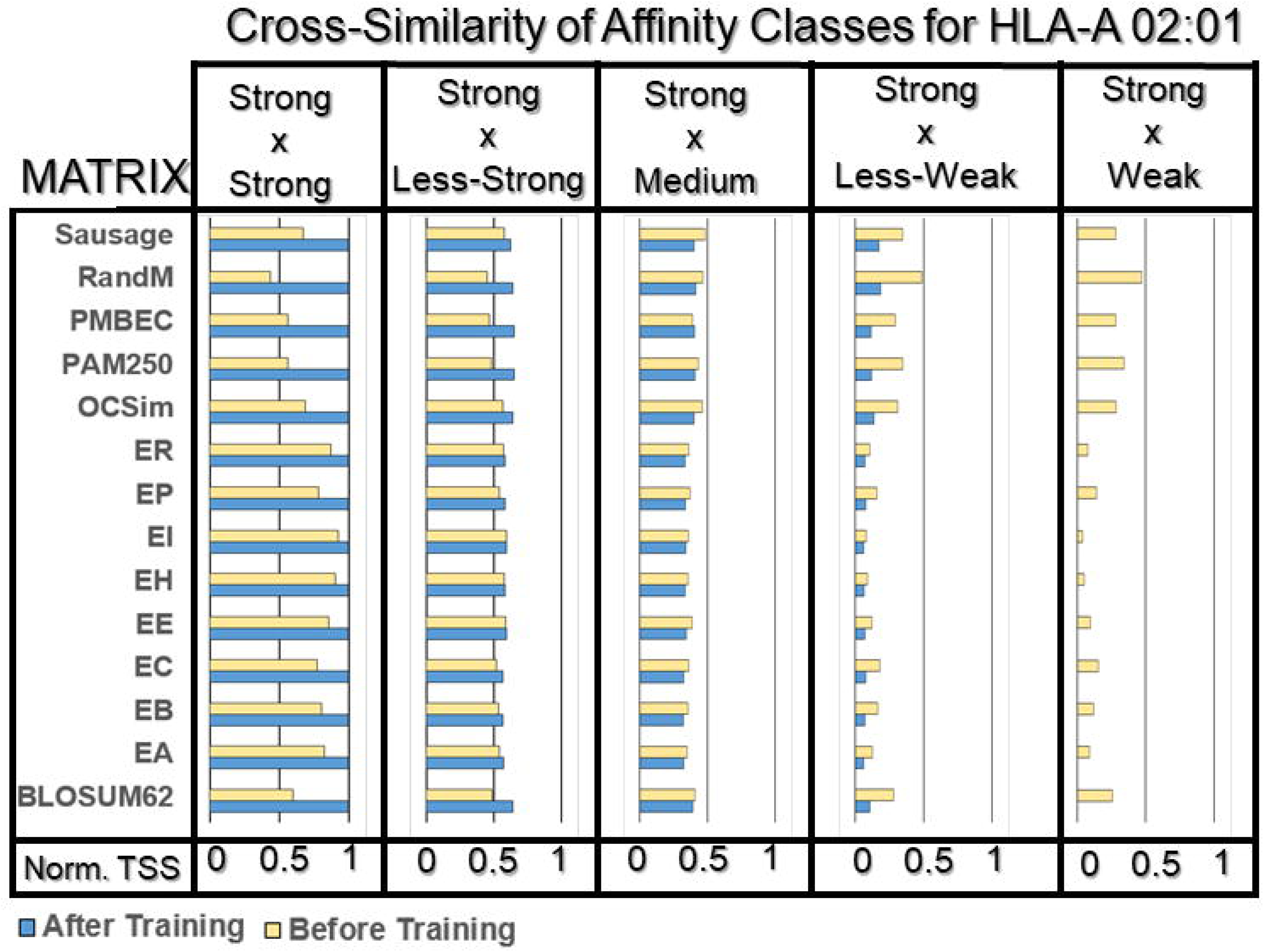
External similarity results for matrices before and after training with *HLA-A 02:01* binders. Five subsets of peptides were created from the full list from each allele. Blue bars represent after training and yellow before training. The TSS for each group to strong binders (TSS_S-LS,M,MW,W_) was calculated in addition to each group to itself (TSS_S-S_, TSS_LS-LS_, TSS_M-M_, TSS_MW-MW_, TSS_W-W_). The y-axis for each bar chart denotes the matrix, the x-axis is the normalized TSS_S-S,LS,M,MW,W_ values. The results show, especially in RandM’s case, that we can improve similarity matrices to predict a trend correlated to binding affinity. This trend is characterized by decreasing TSS_S-S,LS,M,MW,W_ correlating with decreasing binding affinity.

### Correlations with experimental affinity

In the previous work, binding affinity was predicted by placing peptides into semi-quantitative groups of strong, medium and weak by their total similarity score to the strong binding peptide sequences of quartz [25]. The trend of decreasing TSS_Seq-S_ was correlated with experimental affinity by using TSS_Seq-S_ as a threshold to determine whether a peptide would fall into an affinity class (binary classification) [25]. Though significant predictability (70-80%) was obtained using the semi-quantitative scoring method, it falls short of the trend prediction needed to be comparable with MHCFlurry, NetMHC and DeepMHC [12,13,37]. To enable more direct comparisons the Pearson correlation coefficient (linear, Fig 4C) and Spearman rank correlation coefficient (nonlinear, Fig 4B) were calculated, which can determine whether the predicted binding affinity trend (TSS to strong binders) matches the experimental binding affinity trend. In addition, a classifier scheme is included that can recognize whether a peptide is a strong or weak binder by the magnitude of its TSS_Seq-S_. Further, a root mean square error (RMSE) is calculated from the normalized trend of TSS and RMSE to get an idea of close the TSS scores are to the experimental affinity (Fig 4A).

**Fig 4.**
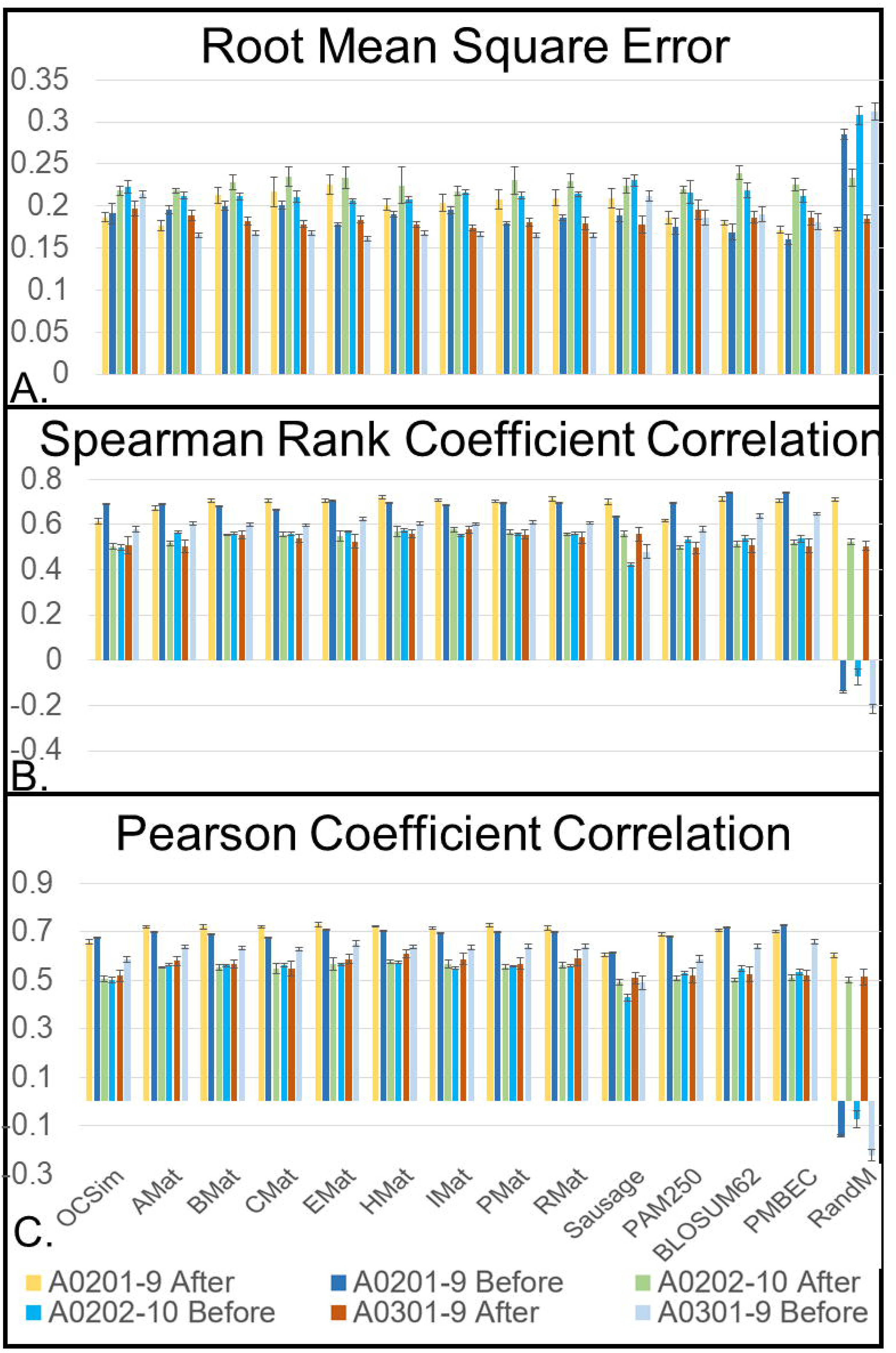
RMSE (A), Spearman Rank (B), and Pearson (C) correlations for TSS trends calculated using trained and untrained matrices. The TSS_Seq-S_ of the *HLA-A*02:01* list was calculated by aligning each peptide with the top binders of the allele and correlating the list of values with the list of experimentally determined binding affinities using linear (Pearson, Fig 4C) and nonlinear (Spearman Rank, Fig 4B) methods. RMSE (Fig 4A) is calculated by obtaining the root mean square of the difference between the normalized (0-1) -pIC_50_ and TSS_Seq-S_. Error bars are 1 standard deviation from the average of each set. The data shows the method can improve literature and calculated matrices, most significantly that a trained randomly initialized matrix (RandM) is more reflective of mutability information in the MHC-I context than all literature and calculated matrices before training.

TSS_Seq-S_ were then correlated with those of the experimental affinity. The values of TSS_Seq-S_ served as the predicted binding affinity ranking and was correlated with the experimentally determined binding affinity using Pearson and Spearman Rank functions. Fig 4 shows the score of each correlation for trained matrices and untrained matrices. The error bars are one standard deviation from the average of these scores. All p-values were less than 0.0005, except for in the case of RandM. Considering the substantially less amount of data used (∼350 peptide sequences for *HLA-A*02:01*) compared with DeepMHC and NetMHC (80% of the full set;∼7200 sequences for *HLA-A*02:01*), the range of 0.5-0.7 is significant and is reflective of mutability information being captured. In addition, the RMSE scores show that in general one TSS score is insufficient to describe the exact binding affinity. While it is clear from the improvement in Pearson and Spearman Rank correlation that these matrices are capturing some similarity information using the method, no matrix alone can produce a TSS ranking exactly correlating with the rest of the set. The integration of several TSS rankings into a single score could prove to be a relevant predictor if they are capturing diverse similarity information unique to their matrix values.

### Binary Classification

Sensitivity and specificity were also recorded as a measure of binary prediction accuracy, shown in Fig 5A and Fig 5B respectively. A binder/nonbinder classification was performed via observing peptides conserved as binders through the magnitude of their TSS_Seq-S_. The sequences having greater than 500 IC_50_ [12] were considered binders. Therefore, peptides given a predicted ranking above the TSS_Seq-S_ threshold correlating with the 500 IC_50_ [12] bar were considered predicted binders. True positives and negatives, and false positive and negatives were calculated by observing which predicted binders were also in the actual binder group.

**Fig 5.**
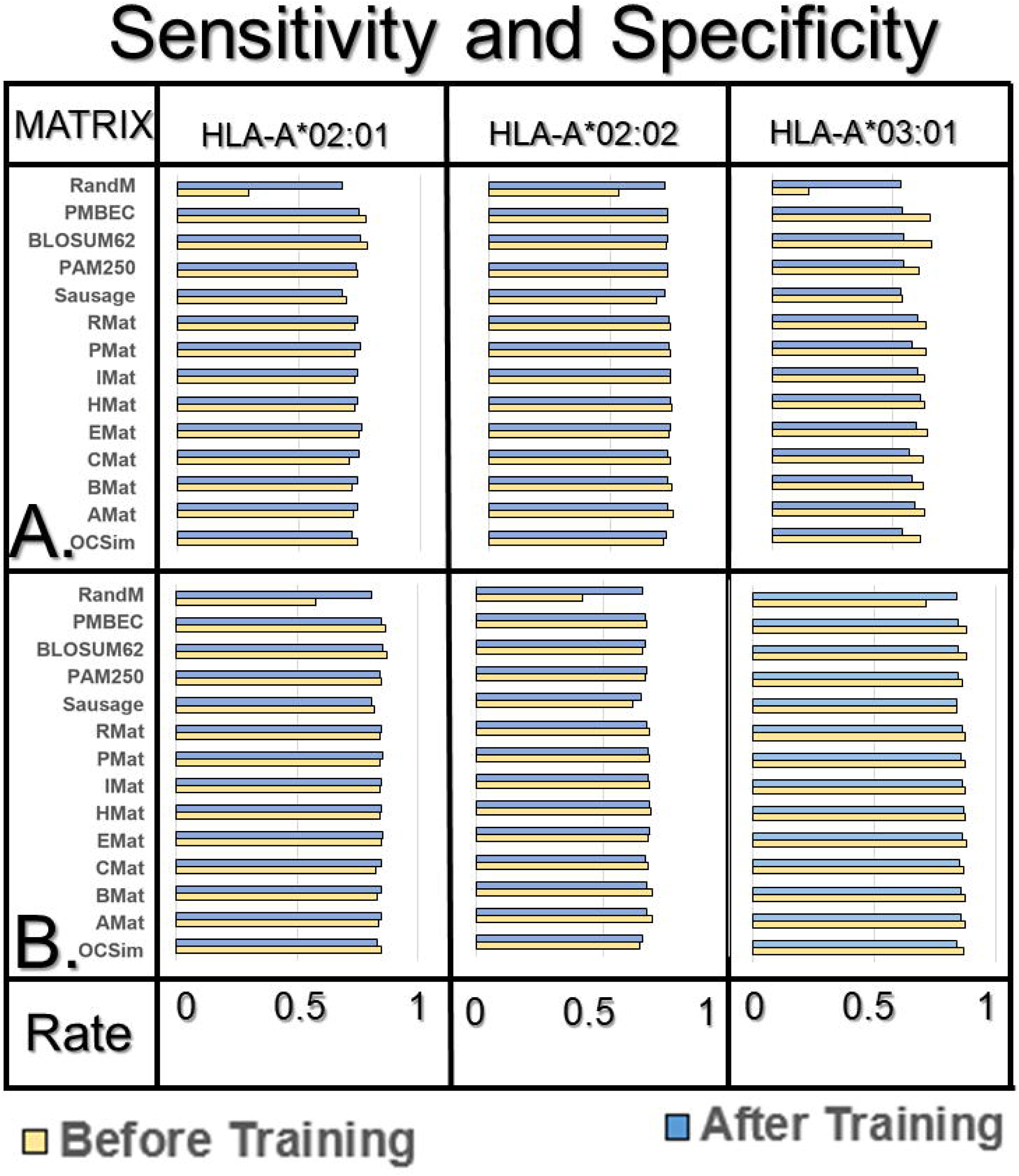
Results of classification analysis. Sensitivity (A) and specificity (B) measures calculated from the results of binary classification of binding. In general, these results show the training function did not improve the predictive ability of any matrix besides RandM, providing evidence that TSS is a relevant predictor while noting the training operation is an ineffective application of TSS.

Across all the matrices a similar specificity/sensitivity was observed before and after training. This indicates the cost function did not improve the ability of the calculated/literature matrices to classify peptides based on TSS values. RandM showed marked improvement across all the alleles but yields lower accuracies than other matrices. This demonstrates that information can be integrated into a similarity matrix up to a limit. In general, the prediction metrics show that the separation of TSS_S-S_ and TSS_S-W_ may not be the appropriate cost function to improve a predictive model. However, TSS_Seq-S_ is a highly relevant predictor of affinity. Though the model was trained on only the dominant features of the peptide set represented by strong binders, the affinity trend was generally conserved by TSS_Seq-S_ scoring.

## Conclusions and Future Work

The predicted correlation range of 0.5-0.7 determined by Pearson and Spearman Rank of the similarity matrix methodology demonstrates similarity matrices can predict functionality (i.e. solid substrate binding specificity) of peptides using the Total Similarity Score. Previous work provided definitive evidence concluding the average similarity score (TSS) of a peptide towards strong binding peptides of an inorganic solid material is positively correlated with the binding affinity of that peptide. Using the Total Similarity Score, we modified a computational method and applied it to a substantially larger dataset to demonstrate that across organic and inorganic materials the metric applies. Though we use substantially lower training data than other methods, similarity matrices were obtained that recognize the dominant features of the strongest binding peptides, which in turn describe those of the weaker binders. Therefore, the strongest binders of the full set can adequately describe the behavior of the remaining peptides Though the training method is insufficient to produce a trend capable of ranking affinity with comparable accuracy to other MHC predictors, we postulate that based on the diversity of the matrices trained that they are capturing different subsections of the total similarity information. Therefore, integrating the trends of multiple matrices into a single score would produce comparable accuracy even when trained on substantially less data. In this work, we show that we can capture similarity information using different matrices and that TSS to strong binders is a relevant predictor of affinity in both organic and inorganic systems.

To uncover the relationship between TSS_Seq-S_ and the experimentally measured affinity, the future work would involve integrating the TSS score with recent statistical learning techniques. If the matrix cannot be optimized, then the value of TSS_Seq-S_ may not be the highest achievable even if the sequence is a strong binder. The sequences with amino acids in similar positions to the strong binding group will, however, tend to give the same average score. Therefore, if the goal is to predict the similarity of sequences based on their positional composition, conserving the common score range will also retain their sequence information. An additional problem may also arise when considering the diversity of the strong binding group. If a given peptide is a strong binder having a completely unique sequence compared to those of the other strong-binding peptides, it will have a low TSS_Seq-S_. TSS scoring assumes that weak and medium binders are mutations of stronger binders. Future methods will capitalize on the information hidden within weak/medium binders and use it to describe the full strong binding space. The full results, gapless Iterative Alignment Python program for calculating similarity matrices, and all the data used to train the matrices are located online on GitHub (https://github.com/Sarikaya-Lab-GEMSEC/Iterative-Alignment-Gapless).

## Acknowledgements

We appreciate the data sets and computational facilities provided GEMSEC labs at the University of Washington.

We declare there were no conflicts of interest for this work.

## Supporting Information Legends

**S1 Fig. Cross-similarity results for the HLA-02:02 allele**. Each bar chart shows the average normalized TSS of the “Strong” affinity class with itself and each other class. The decreasing trend similarity of “Strong” peptides with those of decreasing affinity demonstrates the successful optimization of each matrix for the HLA-A*02:02 allele.

**S2 Fig. Cross-similarity results for the HLA-03:01 allele**. Each bar chart shows the average normalized TSS of the “Strong” affinity class with itself and each other class. The decreasing trend similarity of “Strong” peptides with those of decreasing affinity demonstrates the successful optimization of each matrix for the HLA-A*03:01 allele.

## Notes

#### Summary of Updates

This version of the manuscript has been revised to update several grammatical choices.

https://github.com/Sarikaya-Lab-GEMSEC/Iterative-Alignment-Gapless

