## Supplementary figures and images for "A Generalized Similarity Metric for Predicting Peptide Binding Affinity"

### Supplemental Figure 1

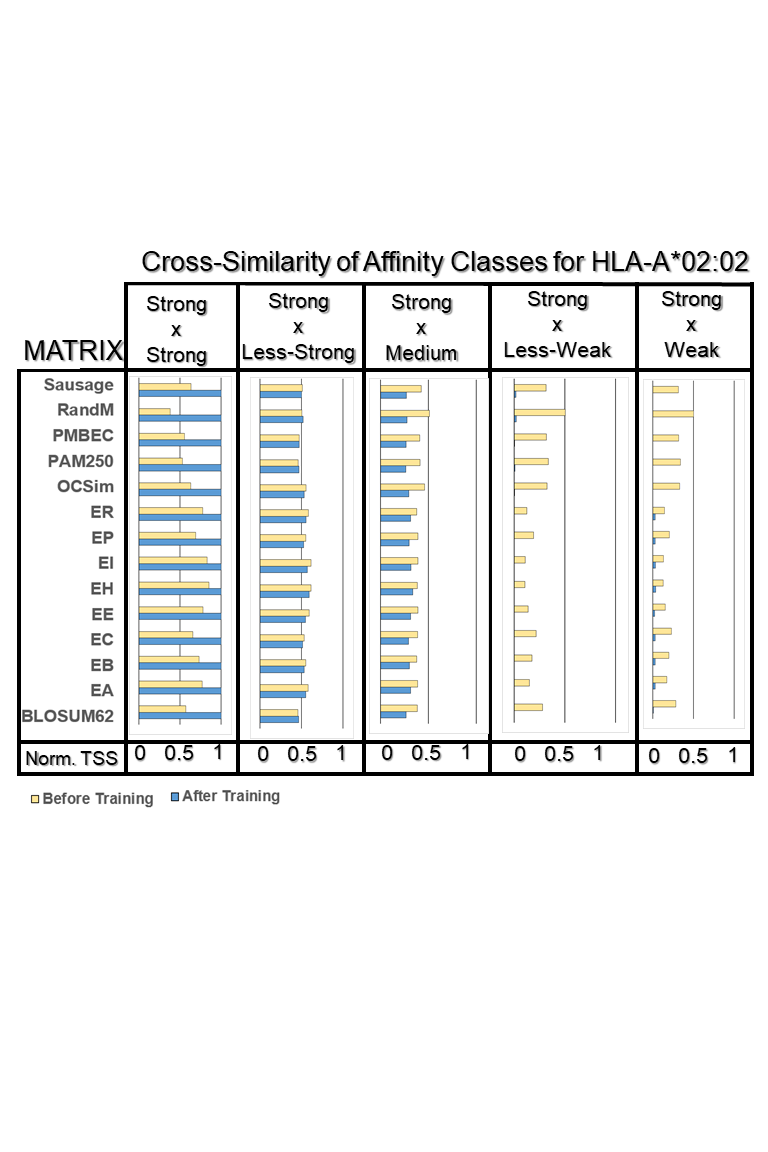

### Supplemental Figure 2

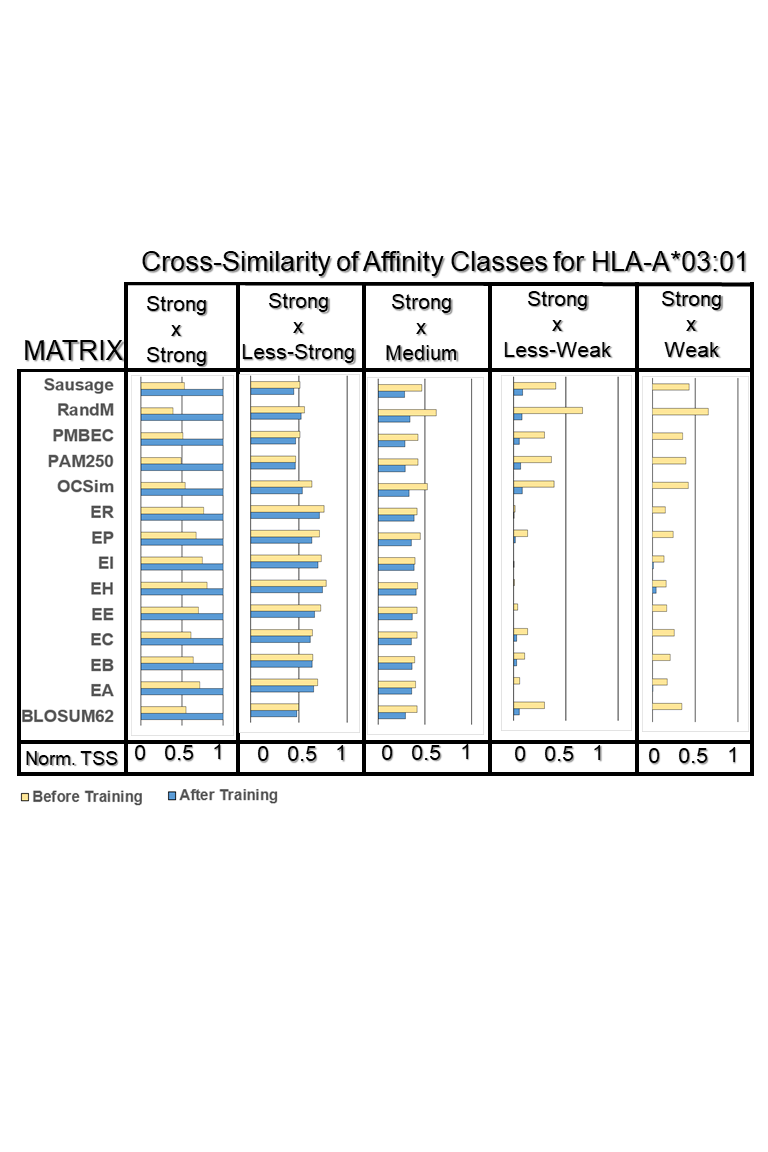
